# *Botryococcus braunii* reduces algal grazing losses to *Daphnia* and *Poterioochromonas* through both chemical and physical interference

**DOI:** 10.1101/2024.02.01.576556

**Authors:** Patrick Thomas, Finn Arn, Micha Freiermuth, Anita Narwani

**Affiliations:** Department of Aquatic Ecology, Swiss Federal Institute of Aquatic Science and Technology (Eawag), Dübendorf, Switzerland; Department of Biology, ETH Zurich, Zurich, Switzerland

**Author notes:** Corresponding author: Patrick Thomas. Swiss Federal Institute of Aquatic Science and Technology (EAWAG), 133 Überlandstasse, 8600 Dübendorf, Switzerland.

**Keywords:** chemical ecology, pest management, synthetic ecology, algal bioproducts, algae milking, allelopathy

## Abstract

Crop protection from algal grazers is a key area of concern, as grazing zooplankton and flagellates can decimate algal crops and impede economic viability of cultivation for biofuels and bioproducts. Inhibition of grazing by chemical and physical interference is one promising solution; however, there have been few empirical tests of this approach that use defense traits innate to algal crop species. Here we conduct an experiment to test whether the hydrocarbon excreting alga *Botryococcus braunii* can mitigate losses to grazing by two distinct grazers, *Daphnia magna* and *Poterioochromonas malhamensis*, due to both chemical inhibition and physical interference linked to large/inedible colonies. We show that chemical and physical defenses interactively reduce the total effect of grazing, thus significantly increasing the biomass of cultures of *B. braunii* and *Nannochloropsis limnetica* when either grazer is present. Specifically, *B. braunii* filtrate alone inhibits grazing and thus weakens top-down control of *N. limnetica* growth rates and final biomass by both grazers; *B. braunii* colonies alone also inhibit biomass losses; and the combination of filtrate and *B. braunii* colonies reveals an interactive effect of both chemical and physical defenses on grazing. Our study demonstrates how community engineering can identify synergies arising from algal cocultivation (e.g., by using industrially relevant strains for crop protection). Such ecological discoveries may help to reduce the costs of large-scale deployment of algal cultivation for sustainable foods, fuels, bioproducts (e.g., bioplastics), and carbon capture.

## 1. Introduction

Protecting microalgal crops from losses to grazers, parasites, and pathogens is a key area of concern, as these organisms can cause rapid pond crashes that threaten the sustainability and economic viability of algae cultivation for bioproducts [1,2]. In general, the most productive, fastest-growing microalgae (i.e., common targets for bioproducts) have small cell sizes which are optimal for maximizing resource uptake and thus growth rates, but are also highly susceptible to grazing by zooplankton including flagellates, rotifers, and cladocerans [3]. Specifically, taxa including *Chlorella, Nannochloropsis*, and *Scenedesmus* are promising due to their high productivity of lipids, proteins, and carbohydrates which can be used for a variety of products (e.g., animal feed, omega-3 fatty acid supplements, biofuels, bioplastics). However, they are also highly susceptible to grazing when grown in unprotected monocultures [2,4]; this can create a trade-off between baseline productivity and grazing resistance capacity. There are several described strategies to overcome this trade-off between maximum productivity and increased risk of pond crash due to grazing or parasitism. Specifically, grazers and fungi can be controlled via external inputs like pesticides and fungicides [5–7] or can be managed by applying pH, ammonia, or salinity changes that inhibit pests but not algae [2,4,8]. However, using one of these solutions in isolation is unlikely to exhibit long-term efficacy against all aquatic grazers and parasites, because biological pest populations can respond adaptively, and their communities are dynamic. This calls for using complementary adaptive management strategies to stabilize large-scale production. Ideally, these interventions will not increase operational costs, energy input, or risk of pesticide release to the environment.

One approach with the potential to stabilize annual productivity is to use designed polycultures, such that different algae species or strains with complementary traits can overcome trade-offs faced by monocultures [9–12], thus simultaneously maintaining baseline productivity and preventing pond crash. This can take a variety of forms, including, e.g., using chemically-defended strains [13] or large-celled/colonial strains [14] to act as an “associational refuge” capable of protecting undefended but faster-growing strains [9]. Despite the promise of using chemically/physically defended strains for crop protection, we currently lack experimental evidence for the efficacy of this approach using industrially relevant strains.

One species of intensive focus in the algae industry is *Botryococcus braunii* due to its high production of extracellular hydrocarbons which can be non-destructively extracted for liquid biofuels (i.e., “algae milking”) [15]; however, it may also have untapped potential in its role for protecting undefended algal crops. Shurin et al. [9] indeed suggested this possibility for crop protection due to *B. braunii*’s ability to chemically inhibit other zooplankton and algae [16], as well as its large colonies offering physical protection from grazing. Therefore, an important step forward in designing robust communities is to test whether *B. braunii* can inhibit grazers without inhibiting undefended target algae (e.g., *Nannochloropsis*).

In this study, we tested whether *B. braunii* inhibits two common and important freshwater grazers and one algal species via both chemical and physical interference. We used *Daphnia magna* and *Poterioochromonas malhamensis* as our two taxonomically and functionally distinct grazer species, and *Nannochloropsis limnetica* as a small (< 4 µm diameter), fast-growing target alga for co-culture with *B. braunii. Daphnia* was chosen as a representative large-bodied (length < 1 mm as juvenile; 1-5 mm as adult), filter-feeding arthropod zooplankton grazer with a larger prey size, while the golden alga *Poterioochromonas* was chosen as a representative small-celled (< 10 µm diameter) flagellate with a smaller prey size that consumes cells via phagocytosis; both are common worldwide and are capable of strong top-down control of algal biomass [2,17]. We experimentally tested the following hypotheses: H1) *B. braunii* reduces algal biomass losses due to grazing via a combination of chemical and physical interference; and H2) *B. braunii* does not chemically inhibit *Nannochloropsis* growth.

## 2. Materials and Methods

### 2.1. Algae and zooplankton cultures

The strain of *B. braunii* used in this experiment was isolated from a bloom in Eawag’s experimental ponds (Dübendorf, Switzerland, 47°24′18.2″ N 8°36′31.7″ E) in April 2023. According to partial 18S sequence matches it belongs to the A-clade and is closely related to previously described strains including, e.g.,OIT-560, OIT-284, and CCAP 807/1. However, it also exhibits traits similar to some B-clade strains of *B. braunii* such as flotation, which is linked to high hydrocarbon content. *Poterioochromonas malhamensis* (CCAP 933/1C) and *Nannochloropsis limnetica* (SAG 18.99) were obtained from culture collections, and *Daphnia magna* was obtained from a clonal culture maintained for several years at Eawag. Prior to the experiment, all cultures were maintained at 20 °C in COMBO medium with double the concentration of all nutrients described in the original medium recipe [18] to allow for higher biomass density (hereafter referred to as 2X COMBO).

### 2.2. Experimental design

For each of the three grazer treatments (grazer-free control, *Poterioochromonas, Daphnia*), there were four *B. braunii* treatments: 1) control (no *B. braunii*); 2) *B. braunii* culture filtrate only, representing only chemical interference; 3) *B. braunii* colonies only, representing primarily physical interference; and 4) *B. braunii* colonies and medium together, representing a combination of both chemical and physical interference of grazing. Treatment 2 (*B. braunii* filtrate only) was made by collecting 0.2-µm filtered medium (i.e., filtrate from Treatment 3) and individually adding 2X COMBO stock nutrients and vitamins to ensure nutrients were initially in excess. Treatment 3 (*B. braunii* colonies) was made by filtering *B. braunii* culture through a 0.2-µm PES Stericup filter and resuspending colonies in fresh 2X COMBO medium. Treatment 4 (colonies and medium) was created by adding 2X COMBO stock nutrients and vitamins directly to a dense culture of *B. braunii*. It should be noted that Treatment 3 (colonies only) isolates the effects of physical interference only earlier in the experiment, while later it likely also represents chemical interference as *B. braunii* metabolites accumulate in the medium. Treatments 2 and 4 are meant to represent the full chemical fingerprint of *B. braunii* medium (i.e., metabolites as well as changes in inorganic chemical parameters such as pH). Aliquots from a stationary phase *Nannochloropsis* culture were added at equal densities to each of the four *B. braunii* treatments (1.3 mL per 50 mL *B. braunii* treatment mixture, a ca. 1% v/v dilution). These mixtures were dispensed into sterile 24-well plates with a volume of 2 mL per well, randomized order in well plates, and 6 replicates per treatment (except for 1 rep. lost in *Poterioochromonas* plate), in Infors incubators set to 20 °C with 12h:12h light:dark cycle and 104 ± 3.2 µmol photons m^-2^ s^-1^. Juvenile *Daphnia* of similar age and size were used for the experiment, with one individual per well; 100 µL of dense *Poterioochromonas* culture was added to each well (i.e., a 5% v/v dilution). Growth was tracked using *in vivo* chlorophyll fluorescence (recorded in terms of relative fluorescence units, RFU) as a proxy for total algal biomass, with excitation/emission at 460/685 nm respectively, using a Biotek® Cytation 5 plate reader. We report algal growth for the first 3 days for *Daphnia* treatments, as *Daphnia* survival was high during this period (96%) reflecting an effective grazing treatment; however, *Daphnia* survival gradually declined after day 3 across all *B. braunii* treatments, thus reducing efficacy of the grazing treatment. We report 9 days of growth for the other grazing treatments. 2-way ANOVAs were used to test for the individual and interactive effects of chemical and physical interference on the growth rate and biomass of all treatments; detailed ANOVA results are in Table S1. Growth rates were calculated as ln(RFU_day3_/RFU_day0_)/3 to capture the initial change in fluorescence during nutrient-replete exponential growth over the first 3 days. We captured images of lugol-preserved cultures for estimating cell densities of each species; however, the large floating colonies of *B. braunii*, as well as flocculation that we observed in *N. limnetica* in treatments containing *B. braunii* medium precluded accurate absolute counts of natural units (cells or colonies). We therefore used in situ chlorophyll-a fluorescence as a proxy for total phytoplankton abundance and report growth rates and grazing-induced losses as changes over time or between grazer and no-grazer controls, respectively. Statistics were performed in R version 4.2.2 [19].

## 3. Results

### 3.1. B. braunii inhibits grazing by Daphnia through both chemical and physical interference

In the control treatment with no *B. braunii* present, *Daphnia* rapidly reduced the *Nannochloropsis* density to a low level (Fig. 1A) over the course of the first three days, with a mean 62% decrease in chl-a fluorescence (Fig. 2A; 74.4% loss if one outlier is excluded). Conversely, the presence of *B. braunii* filtrate inhibited grazing by *Daphnia*, allowing *Nannochloropsis* density to increase despite the presence of the grazers, though chl-a fluorescence was still significantly reduced (mean 29% loss) compared to grazer-free controls. The treatments applying *B. braunii* colonies and colonies + medium provided even further protection from *Daphnia* grazing, with total algal biomass losses of only 4.6% and 10.1%, respectively, compared to grazer-free controls. The 95% CIs overlapping zero indicate no significant declines in biomass (Fig. 2A) or growth rate (Fig. 2C) when *B. braunii* colonies are present. ANOVA revealed significant effects of adding *B. braunii* medium and colonies, as well as a significant interactive effect between the two, in terms of both algae biomass loss (p < 0.02) and growth rate change (p < 0.001); i.e., the effect of adding *B. braunii* medium is reduced when *B. braunii* colonies are present, and vice versa (see Table S1 for ANOVA tables).

**Fig. 1.**
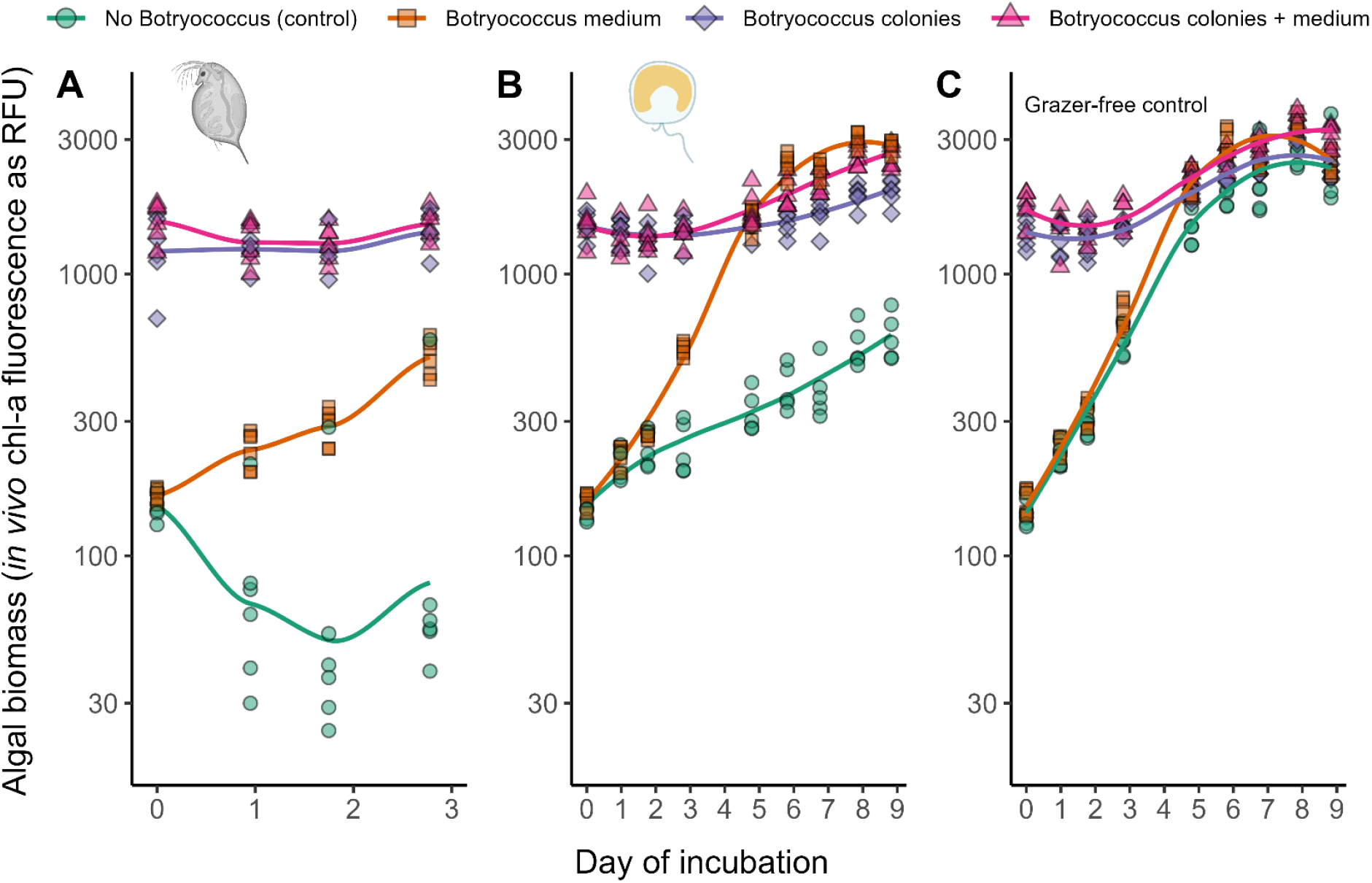
Chlorophyll-a fluorescence (as a proxy for total algal biomass, log-scale) over time in the four *Botryococcus braunii* treatments, with (A) *Daphnia* as grazer, (B) *Poterioochromonas* as grazer, and (C) no grazer added. Both grazers caused strong reductions in biomass (circles in A, B) compared to the grazer-free controls (C) when *Nannochloropsis* was grown alone in fresh medium; however, loss of total algal biomass was mitigated with the addition of *B. braunii*’s filtered culture medium, *B. braunii* colonies alone, and the combination of *B. braunii* medium and colonies. Point show data for individual replicates; lines represent loess fits.

**Fig. 2.**
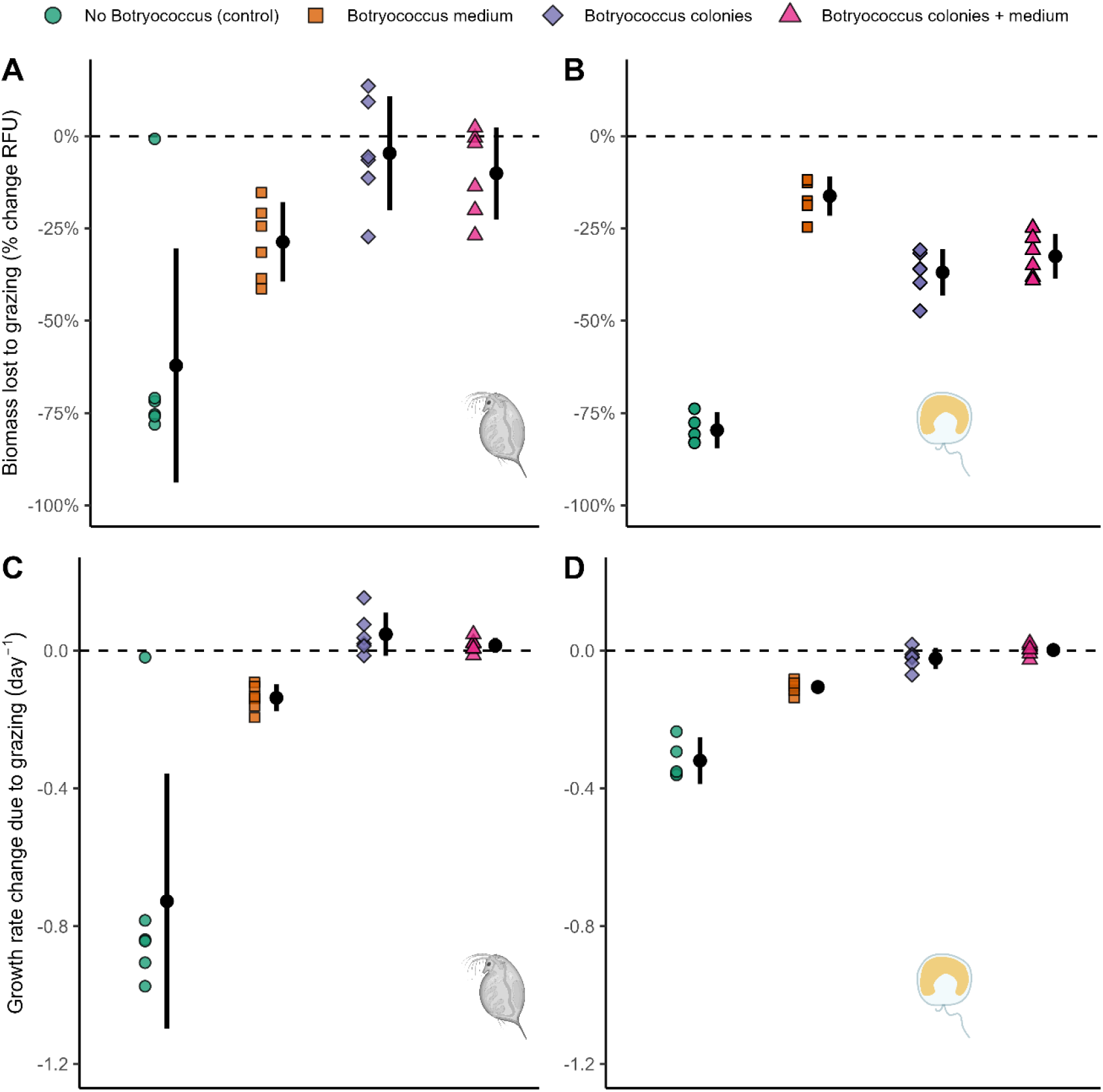
Effects of *B. braunii* treatment on changes in (A, B) maximum chl-a fluorescence (as a proxy for total algal biomass) and (C, D) initial growth rates from day 0-3, due to the presence of grazers (*Daphnia* in A, C; *Poterioochromonas* in B, D). Biomass loss calculations show the % change in fluorescence in each experimental unit relative to the mean grazer-free control in the corresponding *B. braunii* treatment; growth rate changes show the difference in growth rate in each experimental unit compared to the mean grazer-free control. Points are data for individual replicates; black bars show mean and 95% CI per treatment.

### 3.2. B. braunii inhibits grazing by Poterioochromonas through both chemical and physical interference

In the control treatment with no *B. braunii* medium or colonies, *Poterioochromonas* significantly reduced the total algae biomass (mean 80 % decrease in chl-a fluorescence, Fig. 1B, 2B) and growth rates (Fig. 2D) compared to the grazer-free controls. However, the presence of *B. braunii* medium mitigated grazing losses by *Poterioochromonas*, as *Nannochloropsis* chl-a fluorescence only decreased by a mean of 16.2% compared to grazer-free control. In contrast to treatments with *Daphnia* as grazer, the *B. braunii* colonies (37% decrease) and colonies + medium (33 % decrease) treatments allowed for greater grazing losses to *Poterioochromonas* than *B. braunii* medium alone over the full 9-day incubation (Fig. 2B). However, their initial (day 0-3) growth rates did not differ from those of the grazer-free controls (Fig. 2D). ANOVA again yielded a significant interaction term between *B. braunii* medium and colony presence biomass and growth rate (p < 10^−5^ for both; Table S1).1

### 3.3. B. braunii medium enhances Nannochloropsis growth while B. braunii colonies reduce Nannochloropsis growth

Compared to *Nannochloropsis* grown alone in fresh medium (max. fluorescence of 2965 RFU ± 699 SD and growth rate of 0.50 ± 0.01 day^-1^), *Nannochloropsis* grown in the used medium of *B. braunii* had both significantly higher max. fluorescence (3624 RFU ± 371 SD) and initial growth rates (0.54 ± 0.01 day^-1^), indicating an overall enhancement of growth due to *B. braunii* medium (Fig. 1C, Fig. S1). However, presence of *B. braunii* colonies had no overall effect on the maximum fluorescence attained (p = 0.2), indicating that there was not significant overyielding of biomass in co-cultures relative to *Nannochloropsis* monoculture. *B. braunii* colonies had a strong negative effect (p < 0.001) on the initial growth rates of the *Botryococcus*-*Nannochloropsis* co-cultures (which had higher total initial densities) when compared to cultures where only *Nannochloropsis* was present (at low initial densities). These biomass and growth rate results suggest that, as expected, there is niche overlap due to resource competition between the two species. However, this requires further study (i.e., individually tracking *B. braunii* and *Nannochloropsis* cell densities over time, which was impractical in our microcosm study).

## 4. Discussion

The results of our study provide support for our hypotheses that (H1) *B. braunii* inhibits both grazers via a combination of chemical and physical factors, and (H2) *B. braunii* does not chemically inhibit *Nannochloropsis* growth. Our results provide a precedent for engineering algal communities with multiple synergies. Specifically, we propose that *B. braunii* could simultaneously function as a crop protection agent for fast-growing algae such as *Nannochloropsis* that can be harvested for higher-value products, while also producing extracellular hydrocarbons [15].

While we use *Nannochloropsis* as a focal ‘fast-grower’ we imagine that this principle could also apply to other small undefended taxa, e.g., *Synechocystis* that can be used for sustainable PHB bioplastic production [20] or other small strains of interest for feed or fuel. Although in this study we use freshwater organisms, the general principle should apply equally well to brackish or saltwater cultivation. The fact that biomass loss was not completely prevented by *B. braunii* also suggests that this approach should be used in conjunction with complementary approaches [2,4–8,21] to modify the biology and chemistry of pond systems for crop protection to more comprehensively manage complex pond communities. Although the compounds in recycled medium may tend towards overall inhibitory effects [22], there is also potential to identify dissolved metabolites (e.g., plant growth hormones) that stimulate algal growth [23,24].

Although this microcosm study provides a clear proof of concept for a community engineering strategy using *B. braunii* to inhibit grazers and stabilize culture biomass, there are several factors that may determine whether it would be equally effective at industrial scales. For example, we assess only one genotype of each organism; other distinct genotypes of each organism studied here are likely to have differences in key traits (e.g., chemical exudate profiles or colony size across strains/clades of *B. braunii*, differential tolerance to allelopathic inhibition in target algae such as *Nannochloropsis*), which could either enhance or diminish the net protective effect of *B. braunii*. While free fatty acid excretion has been linked to chemical inhibition by *B. braunii* [16,25], there is likely much more to uncover regarding the mechanisms of inhibition caused by the complex suite of bioactive metabolites excreted by different *B. braunii* strains. Different species or genotypes of grazers (or combinations of grazers), and other types of pests like viral, fungal, or bacterial pathogens may (or may not) be inhibited in a similar manner as the two grazers used here. Additionally, effects of other abiotic factors (e.g., temperature, salinity, pH, CO_2_ addition), which are outside the scope of this study, could significantly alter the effects of chemical inhibition and thus be optimized in tandem with the crop protection strategy described in our study. We suggest that an expanding branch of research should explore the potential of chemical and physical inhibition of pests by industrially relevant algae in order to enhance the stability of sustainable bioproduct generation.

## Conclusions

Our study provides a proof of concept that *B. braunii*, an alga that excretes extracellular hydrocarbons, can also protect small, fast-growing algae from grazing pressure, thus providing early evidence for a novel mechanism of protecting algal crop productivity in designed polycultures at mass scale for a suite of complementary renewable bioproducts.

**Supplementary materials:** Table S1 and Fig. S1

**CRediT authorship contribution statement**

**Patrick K. Thomas:** Conceptualization, Methodology, Software, Formal analysis, Investigation, Data curation, Writing – original draft, Visualization. **Finn Arn:** Conceptualization, Methodology, Investigation, Writing – review & editing. **Micha Freiermuth:** Conceptualization, Methodology, Investigation. **Anita Narwani:** Conceptualization, Methodology, Writing – review & editing, Supervision, Funding acquisition.

## Data availability

All data and code used will be publicly available on the Mendeley Data platform as well as Eawag’s institutional repository (https://opendata.eawag.ch/) upon acceptance.

## Acknowledgements

We thank Silvana Käser for maintaining *Daphnia* cultures and for assistance with the *B. braunii* isolation, Marta Reyes for maintaining phytoplankton cultures, Raphaël Bossart for 18S sequencing of *B. braunii*, and Eawag teaching staff for facilitating our collaboration. This manuscript is dedicated to the late Ginger Scales, an omnivorous grazer ca 4.8×10^5^ µm in length who exhibited size-selective feeding on kibble and carrots. This work was funded by a Seed grant from Eawag.

