## Supplementary materials (Table S1, Fig. S1) for "*Botryococcus braunii* reduces algal grazing losses to *Daphnia* and *Poterioochromonas* through both chemical and physical interference"

<u>Contents</u>:

Table S1: detailed ANOVA results

Fig. S1: Effects of *B. braunii* treatment on algal biomass and growth rates with no grazers present

Table S1. 2-way ANOVA results testing effects of *B. braunii* treatment on biomass (max. chl-a fluorescence) and growth rates under different grazer treatments. Significance codes: ***p<0.001, **p<0.01, *p<0.05.

**(A) Effects of Botryococcus treatment on chl-a biomass change with Daphnia as grazer**

| | Df | Sum of squares | Mean squares | F statistic | p value | significance |
|---|---|---|---|---|---|---|
| bot_medium | 1 | 1180 | 1180 | 3.434 | 0.0787 | . |
| bot_colonies | 1 | 8678 | 8678 | 25.259 | 6.48E-05 | *** |
| bot_medium:bot_colonies | 1 | 2272 | 2272 | 6.614 | 0.0182 | * |
| Residuals | 20 | 6871 | 344 | | | |

**(B) Effects of Botryococcus treatment on growth rate change with Daphnia as grazer**

| | Df | Sum of squares | Mean squares | F statistic | p value | significance |
|---|---|---|---|---|---|---|
| bot_medium | 1 | 0.4662 | 0.4662 | 14.35 | 0.001153 | ** |
| bot_colonies | 1 | 1.2888 | 1.2888 | 39.68 | 3.78E-06 | *** |
| bot_medium:bot_colonies | 1 | 0.5816 | 0.5816 | 17.91 | 0.000409 | *** |
| Residuals | 20 | 0.6496 | 0.0325 | | | |

**(C) Effects of Botryococcus treatment on chl-a biomass change with Poterioochromonas as grazer**

| | Df | Sum of squares | Mean squares | F statistic | p value | significance |
|---|---|---|---|---|---|---|
| bot_medium | 1 | 5854 | 5854 | 205.3 | 1.23E-11 | *** |
| bot_colonies | 1 | 796 | 796 | 27.9 | 4.25E-05 | *** |
| bot_medium:bot_colonies | 1 | 4985 | 4985 | 174.8 | 4.95E-11 | *** |
| Residuals | 19 | 542 | 29 | | | |

**(D) Effects of Botryococcus treatment on growth rate change with Poterioochromonas as grazer**

| | Df | Sum of squares | Mean squares | F statistic | p value | significance |
|---|---|---|---|---|---|---|
| bot_medium | 1 | 0.06428 | 0.06428 | 62.63 | 1.97E-07 | *** |
| bot_colonies | 1 | 0.22312 | 0.22312 | 217.39 | 7.45E-12 | *** |
| bot_medium:bot_colonies | 1 | 0.05086 | 0.05086 | 49.55 | 1.06E-06 | *** |
| Residuals | 19 | 0.0195 | 0.00103 | | | |

**(E) Effects of Botryococcus treatment on chl-a biomass with grazers absent**

| | Df | Sum of squares | Mean squares | F statistic | p value | significance |
|---|---|---|---|---|---|---|
| bot_medium | 1 | 0.2985 | 0.29849 | 17.074 | 0.000517 | *** |
| bot_colonies | 1 | 0.0319 | 0.03195 | 1.828 | 0.191512 | |
| bot_medium:bot_colonies | 1 | 0.0002 | 0.00023 | 0.013 | 0.910388 | |
| Residuals | 20 | 0.3496 | 0.01748 | | | |

**(F) Effects of Botryococcus treatment on growth rate change with grazers absent**

| | Df | Sum of squares | Mean squares | F statistic | p value | significance |
|---|---|---|---|---|---|---|
| bot_medium | 1 | 0.0002 | 0.0002 | 0.529 | 0.475547 | |
| bot_colonies | 1 | 1.6855 | 1.6855 | 4244.987 | <2E-16 | *** |
| bot_medium:bot_colonies | 1 | 0.0082 | 0.0082 | 20.694 | 0.000195 | *** |
| Residuals | 20 | 0.0079 | 0.0004 | | | |

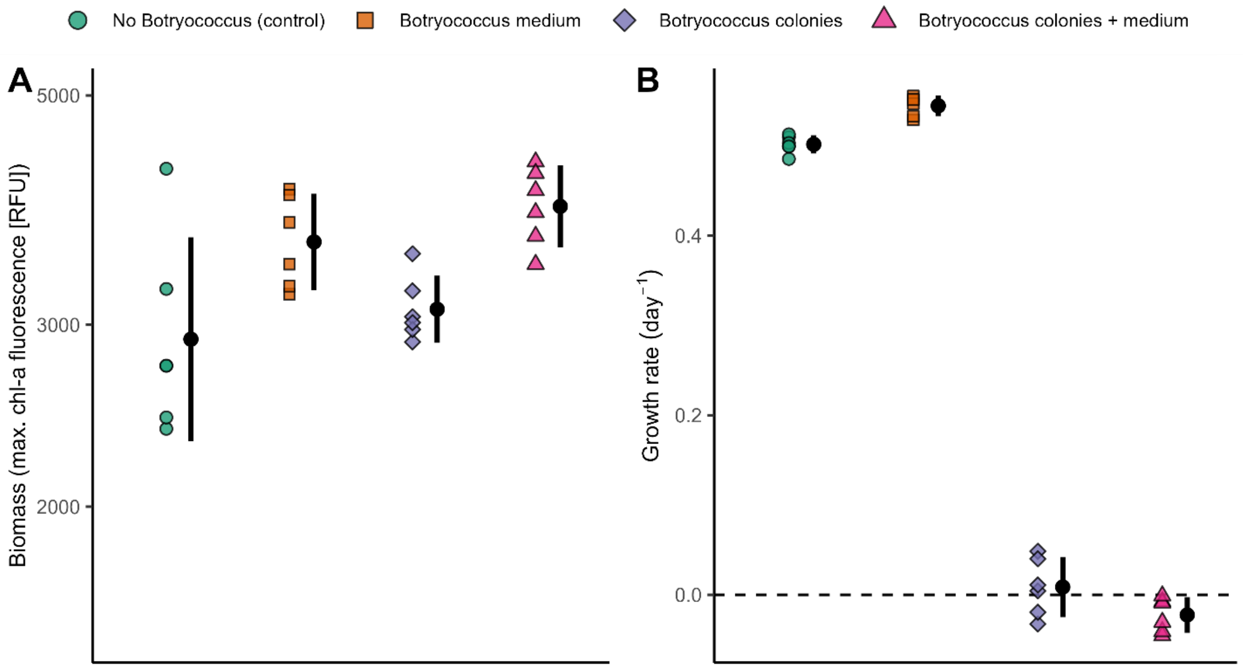


Fig. S1. Biomass in terms of maximum chl-a fluorescence (A) and initial day 0-3 growth rates (B) for different *B. braunii* treatments when no grazer is present. Points are data for individual replicates; black bars show mean and 95% CI per treatment.
